# Increased cerebrospinal fluid and plasma apoE glycosylation is associated with reduced levels of Alzheimer’s disease biomarkers

**DOI:** 10.1101/2024.12.20.629619

**Authors:** Dobrin Nedelkov, Zoe E. Tsokolas, Matheus Scarpatto Rodrigues, Isabel Sible, S. Duke Han, Bilal E. Kerman, Michael Renteln, Wendy J. Mack, Tharick A. Pascoal, Hussein N Yassine, the Alzheimer’s Disease Neuroimaging Initiative

## Abstract

The apolipoprotein E (*APOE*) ε4 allele is the strongest genetic risk factor for Alzheimer’s disease (AD). ApoE is glycosylated with an O-linked Core-1 sialylated glycan at several sites, yet the impact and function of this glycosylation on AD biomarkers remains unclear. We examined apoE glycosylation in a cohort of cerebrospinal fluid (CSF, n=181) and plasma (n= 178) samples from the Alzheimer’s Disease Neuroimaging Initiative (ADNI) stratified into 4 groups: cognitively normal (CN), Mild Cognitive Impairment (MCI), progressors and non-progressors based on delayed word recall performance over 4 years. We observed decreasing glycosylation from apoE2 > apoE3 > apoE4 in CSF, and in plasma (apoE3 > apoE4). ApoE glycosylation was reduced in the MCI compared with CN groups, and in progressors compared to non-progressors. In CSF, higher apoE glycosylation associated cross-sectionally with lower total tau (t-tau), p-tau181, and with higher Aβ_1-42_. Similar associations of apoE glycosylation with higher Aβ_1-42_ were observed in plasma. In CSF, greater apoE4 glycosylation was associated with lower t-tau and p-tau181. Over a 6-year period, higher baseline levels of CSF apoE glycosylation predicted lower rates of increase in CSF t-tau and p-tau181 and lower rates of decrease in CSF Aβ_1-42_. These results indicate strong associations of apoE glycosylation with biomarkers of AD pathology independent of apoE genotype, warranting a deeper understanding of the functional role of apoE glycosylation on AD tau pathology.

## Introduction

The human apolipoprotein E gene (*APOE*) contains three major polymorphic alleles with distinct distribution across ethnicity and geography (ε2: 5-10%, ε3: 65-70%, and ε4: 15-25%) [1]. The ε4 allele is the strongest genetic risk factor for Alzheimer’s disease (AD), with its frequency increased to ∼ 40% in patients with AD [2]. Having one copy of the ε4 allele increases the risk for late-onset AD three-fold, while individuals with two copies of ε4 have an eight- to twelve-fold increased risk of developing AD [2–4]. The three distinct APOE alleles translate into three apoE isoforms that differ from each other at amino acid positions 112 and 158: apoE2 (Cys112, Cys158), apoE3 (Cys112, Arg158), and apoE4 (Arg112, Arg158). The three apoE isoforms bind differently to lipids, receptors, and amyloid-β (Aβ) [5–11], but the exact mechanism by which the isoforms contribute individually to AD pathogenesis is not completely understood.

ApoE is an already-glycosylated protein with an O-linked N-acetylgalactosamine- Galactose disaccharide (-GalNAc-Gal); this Core-1 glycan (also known as T-antigen) can be further sialylated with 1 or 2 sialic acids (Sia) [12]. The primary site of apoE glycosylation is at Thr194, which resides on the hinge region of the apoE three-dimensional structure [13]. Ser290 near the C-terminus was also identified as a major glycan attachment site [14]. More recent mass spectrometry-based studies delineated several additional glycosylation sites, including Thr8, Thr18, Ser197, Ser263, Thr289, and Ser296 [15–18]. Post-translational glycosylation of apoE may mediate its intermolecular interactions and enhance its solubility and stability. Glycosylation may also protect against self-association, spontaneous aggregation, and fibril formation in AD and atherosclerotic plaques [14]. In a Niemann-Pick Type C disease animal model, sialic acid residues were detected on neuronal apoE, indicating a change in glycosylation which correlated with increased Aβ_1-42_ before neurological abnormalities appear [19]. The sialic acids at the end of the apoE glycan may contribute to interaction with Aβ [20], mediate/stabilize intermolecular interactions with the polar phospholipid heads of micelle [14], and are needed for binding to HDL [21, 22]. A larger proportion of apoE is glycosylated and sialylated in CSF than in plasma [23–26], but the degree to which apoE glycosylation in CSF affects AD pathogenesis remains unclear.

We have recently developed an assay that detects all apoE isoforms and glycoforms in a manner that preserves isoform-specific glycosylation [27]. Using this assay on a small cohort of matched plasma and CSF samples (n=22), we discovered for the first time a trend of decreasing glycosylation of apoE in CSF from apoE2 > apoE3 > apoE4, with apoE4 being the least glycosylated isoform [27]. This trend and findings were recently validated with a larger cohort (n=106) [28]. Here, we seek to further examine the relationship of apoE glycosylation with AD biomarkers including total tau (t-tau), phosphorylated tau 181 (p-tau181), and Aβ_1-42_, as well as with the cognitive performance of individuals at risk for or with AD, using a larger cohort of plasma and CSF samples with longitudinal biomarker data.

## Materials and Methods

### Study Population

Data and samples used in the preparation of this article were obtained from the Alzheimer’s Disease Neuroimaging Initiative (ADNI) database (adni.loni.usc.edu). The ADNI was launched in 2003 as a public-private partnership, led by Principal Investigator Michael W. Weiner, MD. The primary goal of ADNI has been to test whether serial magnetic resonance imaging (MRI), positron emission tomography (PET), other biological markers, and clinical and neuropsychological assessment can be combined to measure the progression of mild cognitive impairment (MCI) and early AD.

From the ADNI dataset, our analysis included individuals with available medical data, clinical assessment, and measures of post-translational glycosylation of apoE in both CSF (n = 181 participants) and plasma 178 (n = 178 participants). We stratified individuals in groups based on the participants’ delayed recall test performance as assessed by the Rey Auditory Verbal Learning Test (RAVLT, Rey, 1964) at baseline and *APOE* genotype (**Table 1**). Therefore, the cohort was stratified into four clinical groups: 1) cognitively normal (CN) (n=41) stable group, with participants having no cognitive impairment (at baseline assessment) and <10% RAVLT decline between baseline visit and 48-months follow-up; 2) CN decline group (n=43), with participants having no cognitive impairment (at baseline assessment) and >10% RAVLT decline between baseline and 48-months follow up; 3) Late Mild Cognitive Impairment (LMCI) stable group (n=48), with participants having mild cognitive impairment at baseline (MCI) and without AD progression between baseline and 48-months; and 4) LMCI decline group (n=49), with participants having MCI at baseline with AD progression between baseline and 48-months.

**Table 1.** Study Cohort’s Demographic and Clinical Characteristics.

| **Table 1. Cohort characteristics** |  |  |  |  |  |
| --- | --- | --- | --- | --- | --- |
|  | **Whole Cohort** | **Cognitively Normal Non-Progressors** | **Cognitively Normal Progressors** | **Mild Cognitive Impairment Non-Progressors** | **Mild Cognitive Impairment Progressors** |
| ***N* (at baseline)** | 181 | 41 | 43 | 48 | 49 |
| **Sex (*N*, % Male)** | 103 (56.9%) | 22 (53.7%) | 20 (46.5%) | 31 (64.6%) | 30 (61.2%) |
| **Age (Mean ± SD)** | 73.3 ± 6.4 | 75.0 ± 5.6 | 74.8 ± 4.6 | 71.7 ± 6.9 | 72.3 ± 7.0 |
| **Years of Education (Mean ± SD)** | 16.2 ± 2.7 | 15.6 ± 2.8 | 16.3 ± 2.8 | 16.4 ± 2.8 | 16.0 ± 2.7 |
| **Race (*N*, %)** |  |  |  |  |  |
| Asian/Pacific Islander | 10 (9.9%) | 3 (7.3%) | 1 (2.3%) | 3 (6.3%) | 3 (6.1%) |
| Caucasian | 170 (93.9%) | 38 (92.7%) | 42 (97.7%) | 45 (93.8%) | 45 (91.8%) |
| Other/Unknown | 1 (0.5%) | 0 (0%) | 0 (0%) | 0 (0%) | 1 (2.0%) |
| ***APOE* Genotype (*N*, %)** |  |  |  |  |  |
| E2/E3 | 12 (6.6%) | 4 (9.8%) | 3 (7.0%) | 3 (6.3%) | 2 (4.1%) |
| E3/E3 | 82 (45.3%) | 17 (41.5%) | 21 (48.8%) | 22 (45.8%) | 22 (44.9%) |
| E3/E4 | 56 (30.9%) | 15 (36.6%) | 14 (32.6%) | 14 (29.2%) | 13 (26.5%) |
| E4/E4 | 31 (17.1%) | 5 (12.2%) | 5 (11.6%) | 9 (18.8%) | 12 (24.5%) |
| ***Clinical***<br>***Dementia Rating***<br>***Score (N, %)*** |  |  |  |  |  |
| 0-0.5 | 102 (56.4%) | 41 (100%) | 43 (100%) | 9 (18.8%) | 9 (18.4%) |
| 1-1.5 | 51 (28.2%) | 0 (0%) | 0 (0%) | 24 (50.0%) | 27 (55.1%) |
| 2-2.5 | 19 (10.5%) | 0 (0%) | 0 (0%) | 10 (20.8%) | 9 (18.4%) |
| 3+ | 9 (5.0%) | 0 (0%) | 0 (0%) | 5 (10.4%) | 4 (8.2%) |
| ***CSF Biomarkers***<br>***(Mean ± SD)*** |  |  |  |  |  |
| Aβ1-42<br>(pg/mL) | 980.1 ± 462.8 | 1089.9 ± 443.6 | 1058.2 ± 494.6 | 915.9 ± 483.0 | 869.4 ± 404.3 |
| Total tau<br>(pg/mL) | 287.1 ± 113.2 | 268.8 ± 78.7 | 297.7 ± 108.7 | 277.7 ± 97.1 | 307.6 ± 149.9 |
| p-tau181<br>(pg/mL) | 27.4 ± 12.5 | 25.2 ± 8.7 | 28.9 ± 12.4 | 26.3 ± 10.5 | 29.6 ± 16.5 |

### Study Data

Biomarker data were available from CSF samples collected during annual visits spanning a 6-year period. The concentrations of Aβ1-42, t-tau, and p-tau181 in CSF were measured on the Luminex platform INNO-BIA AlzBio3 RUO test, using methods of micro-bead-based multiplex immunoassay as previously described [29]. For cognitive measures, the RAVLT scores accuracy of delayed memory recall by % forgotten on a list-recall paradigm was used [30]. RAVLT cognitive performance test results were available from each annual visit over the 6-year period.

### Glycosylation Measurement

Plasma and CSF samples from the first (baseline) visit were analyzed the apoE glycosylation using ApoE mass spectrometric immunoassay (MSIA), as described previously [27]. The MSIA detects each of the three ApoE isoforms and their glycoforms at the intact protein level, even in samples from heterozygous individuals. In short, apoE was affinity captured from plasma and CSF using MSIA tips that were derivatized with ApoE-specific antibody. MALDI-TOF mass spectrometer (Autoflex III MALDI-TOF, Bruker, Billerica, MA) was used to analyze the eluted intact ApoE in a positive ion mode, with a mass range 7 to 70 kDa, 700 ns delay, 20.00 kV and 18.45 kV ion source voltages, and < 7000 Da signal suppression. Mass spectra were baseline subtracted (Convex Hull algorithm with 0.8 flatness) and smoothed (Savizky Golay algorithm with 5 m/z width and 1 cycle) using the Bruker Flex Analysis software. Zebra 1.0 software (Intrinsic Bioprobes Inc.) was used to quantify and tabulate peak intensities of the *APOE* isoforms and glycoforms. To obtain glycosylation percentages, glycoform peak intensities for each isoform were summed and divided by the sum of all peak intensities (glycosylated and unglycosylated) for that isoform. Similarly, to obtain the secondary glycosylation percentage (the additional glycosylation of an already singly glycosylated apoE at a site that is different from the primary glycosylation site), only the peak intensities representing secondary glycosylation were summed and divided by the sum of all peak intensities for that isoform.

### Statistical Analysis

Cross sectional statistical analyses were performed using GraphPad Prism 7 and R (version 4.3.3). Data normality was assessed by the Shapiro-Wilk test. Biomarker concentrations were standardized to a mean of 0 and SD of 1 and used in cross-sectional and longitudinal analyses to improve model stability. To evaluate for differences in the overall apoE glycosylation between plasma and CSF, unpaired t-test was performed at a 5% false discovery rate. To identify differences among the glycosylation of the *APOE*2, *APOE*3, and *APOE*4 isoforms, one-way ANOVA with Tukey multiple comparisons test was performed for normally distributed data, and Kruskal-Wallis test with Dunn’s multiple comparisons test was performed for non-normally distributed data. Linear regression models adjusted by age, sex, race, education, *APOE*4, and clinical group were conducted to examine associations between apoE glycosylation and AD biomarkers. Statistical differences were assessed via Pearson’s correlation (for normally distributed measures) or Spearman’s rank correlation (for non-normally distributed measures). We further stratified the total cohort into two groups based on cognitive progression as determined by the % forgetting on the RAVLT score (non-progressors: CN stable and LMCI stable; and progressors: CN- decline and LMCI-decline) to examine differences across total and secondary apoE glycosylation levels. Longitudinal statistical analyses were performed using R (version 4.3.3).

Associations between baseline assessments of CSF and plasma apoE glycosylation and longitudinal assessments of CSF t-tau, p-tau181, Aβ_1-42_, and cognitive performance over a six-year period were tested using linear mixed effects models with the lme4 R package (version 1.1-35.2). The repeatedly measured dependent variables were t-tau, p-tau181, Aβ_1-42_, and cognitive performance as assessed by RAVLT (measured annually for a duration of 6 years). The primary fixed effects of interest were tertile categories of the independent apoE glycosylation variable. Additional covariates were age, sex, race, education, clinical group, *APOE*4, data collection site, and year of test visit (relative to baseline, with year = 0 at baseline visit). To account for random variation across testing locations and within-participant correlations due to repeated measures, study site was modeled as a random effect; a subject-level random intercept was modeled to account for correlated outcome data due to repeated measures. Time (year)-by-tertile interactions were included in longitudinal modeling to test for differences in rates of change in the dependent measure over time by tertiles of glycosylation. We modeled these glycosylation categories as the primary independent variables of interest and adjusted for clinical group, APOE4, age, sex, race, education, study site, and years from baseline assessment. We further adjusted for participant ID to account for within-subject variability, as a random effect. To categorize tertiles, baseline glycosylation samples were divided into three groups based on abundance levels of total and secondary glycosylation (1st tertile = lowest one third % of glycosylation; 2nd = middle; 3rd = highest degree of glycosylation). To test differences in rates of change over time by glycosylation tertiles, we included a tertile-by-years interaction in our linear models.

## Results

### Cross-sectional analyses

#### Higher CSF APOE glycosylation is associated with lower AD pathology and better cognitive performance

ApoE glycosylation levels from baseline collection were first examined as a function of the individual *APOE* isoforms, as our previous work showed a trend of decreasing glycosylation from *APOE2* > *APOE3* > *APOE4* [27, 28]. Here we observe the same trend of decreasing glycosylation: for CSF total apoE glycosylation, the mean (SD) percentage glycosylation was 74.5% (2.15%) for *APOE*2, 71.1% (4.27%) for *APOE*3, and 67.5% (4.32%) for *APOE*4 (ANOVA p < 0.0001), with statistically significant differences for glycosylated *APOE*2 vs. *APOE*3 (p=0.02), *APOE*2 vs. *APOE*4 (p < 0.0001), and *APOE*3 vs. *APOE*4 (p < 0.0001) (**Fig. 1a**). For the CSF secondary APOE glycosylation, the mean (SD) percentage glycosylation was 23.3% (2.4%) for *APOE*2, 20.9% (3.71%) for *APOE*3, and 15.6% (2.77%) for *APOE*4 (ANOVA p < 0.0001), with statistically significant differences observed for *APOE*4 vs. *APOE*2 (p < 0.0001) and *APOE*4 vs. *APOE*3 (p < 0.0001) (**Fig. 1b**). In plasma, the mean (SD) total percentage APOE glycosylation was 11.9% (3.53%) for *APOE*2, 12.8% (3.96%) for *APOE*3, and 9.70% (3.01%) for *APOE*4 (ANOVA p < 0.0001), with statistically significant differences between *APOE*4 and *APOE*3 (p < 0.0001) (**Fig. 1c**). Overall, the total percentage glycosylation of apoE in CSF was 69.2% (4.11%), much higher than the 11.8% (3.88%) total glycosylation observed in plasma (p < 0.0001) (**Fig. 1 Supplement**), which is in close agreement with our previous results [27, 28].

**Fig. 1.**
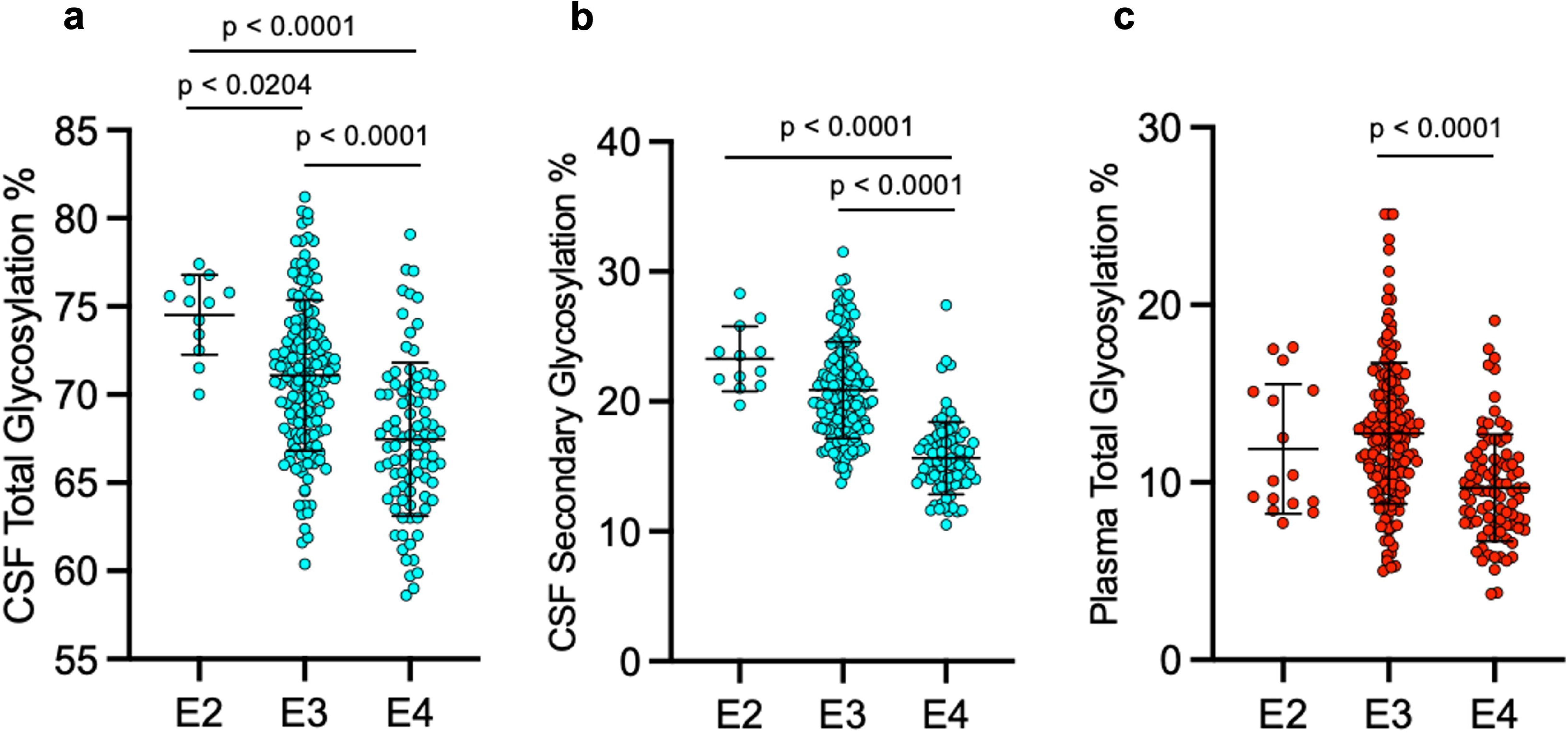
ApoE glycosylation percentage by isoform (APOE2, APOE3, and APOE4). **a)** CSF total apoE glycosylation, **b)** CSF secondary apoE glycosylation, and **c)** Plasma apoE glycosylation. Groups were compared using ANOVA.

We conducted independent two-sample t-tests to compare differences in mean apoE glycosylation across each of the 4 clinical groups and used the Bonferroni method to correct for multiple comparisons. In the CN and LMCI groups, the mean (SD) percentage CSF secondary apoE glycosylation of 19.5% (4.03%) in the CN group was significantly higher than the mean of 18.0% (3.32%) in the LMCI group (p=0.0085) (**Fig. 2a**). The total cohort was also split into two groups based on memory performance as determined by the RAVLT score (non-progressors; progressors). The mean percentage CSF secondary apoE glycosylation of 19.3% (3.38%) in the overall non-progressors group was significantly higher than the mean of 18.1% (3.98%) in the overall progressors group (p = 0.022) (**Fig. 2b**). This difference was primarily driven by the CN group, with CN non-progressors having significantly higher secondary apoE glycosylation levels than CN progressors (p = 0.012) (**Fig. 2c**). However, MCI progressors did not differ from non-progressors (p=0.5) (**Figure 2d**).

**Fig. 2.**
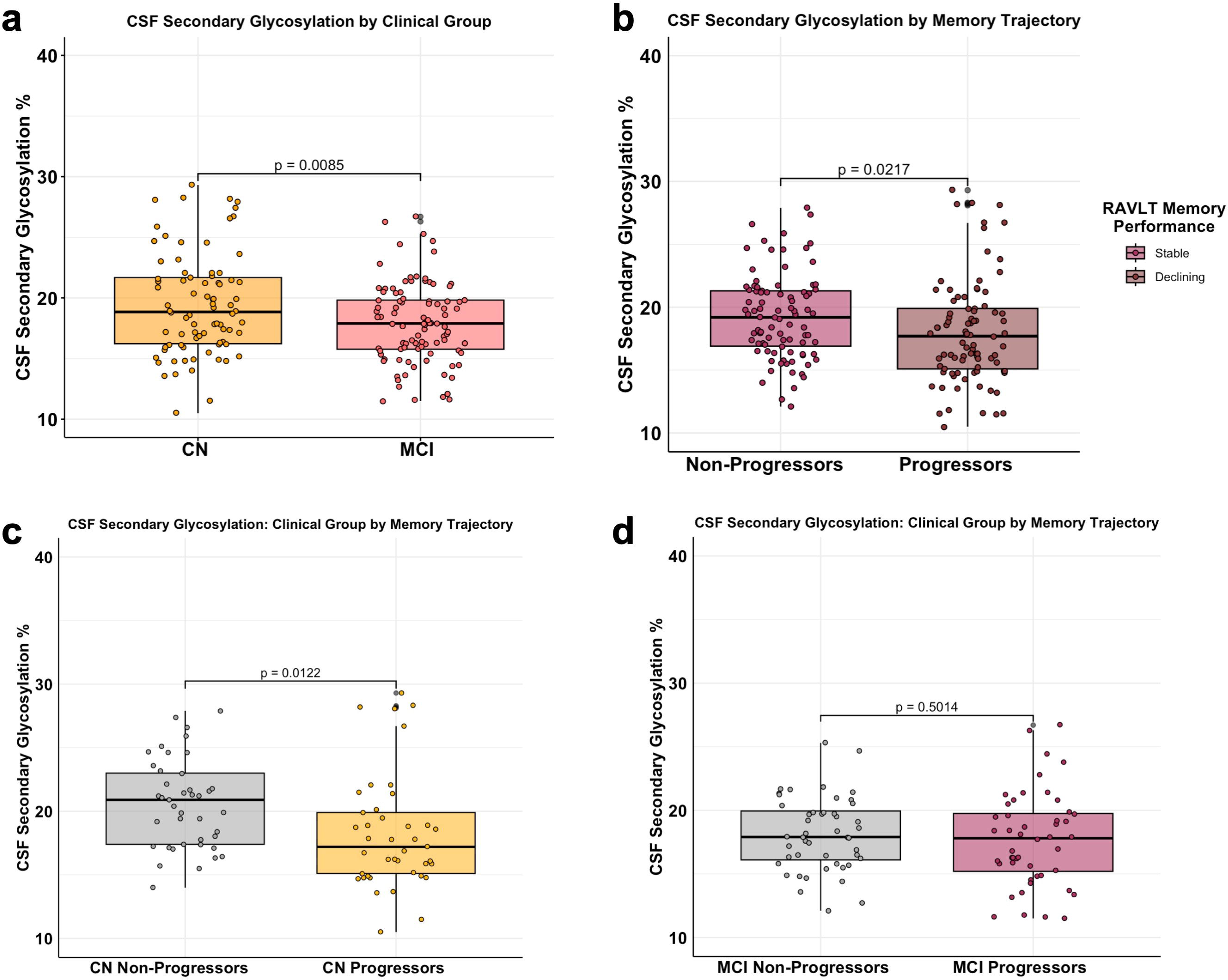
CSF apoE secondary glycosylation at baseline in CN and LMCI groups, and in non-progressors and progressors. Independent two-sample t-tests were performed to compare group differences in CSF apoE secondary glycosylation.

At baseline, higher CSF total apoE glycosylation was significantly associated with lower total CSF tau (r = -0.17; p = 0.016) (**Fig. 3a**) but not Aβ_1-42_ (r = -0.06; p = 0.58) (**Fig. 3b**), ptau-181 (r = -0.12; p = 0.068) (**Fig. 3c**), or RAVLT (r = -0.03; p = 0.63) (**Fig. 3d**). Higher CSF secondary apoE glycosylation was not associated with lower total CSF tau (r = -0.04; p = 0.3) (**Fig. 2a Supplement**), CSF Aβ_1-42_ (r = -0.1; p = 0.32), (**Fig. 2b Supplement**), CSF ptau-181 (r = 0.01; p = 0.62) (**Fig. 2c Supplement**), or RAVLT (r = 0.05; p = 0.41) (**Fig. 2d Supplement**). For plasma, higher total apoE glycosylation was significantly associated with lower Aβ_1-42_ (r = -0.08; p = 0.0006) (**Fig. 3c Supplement**).

**Fig. 3.**
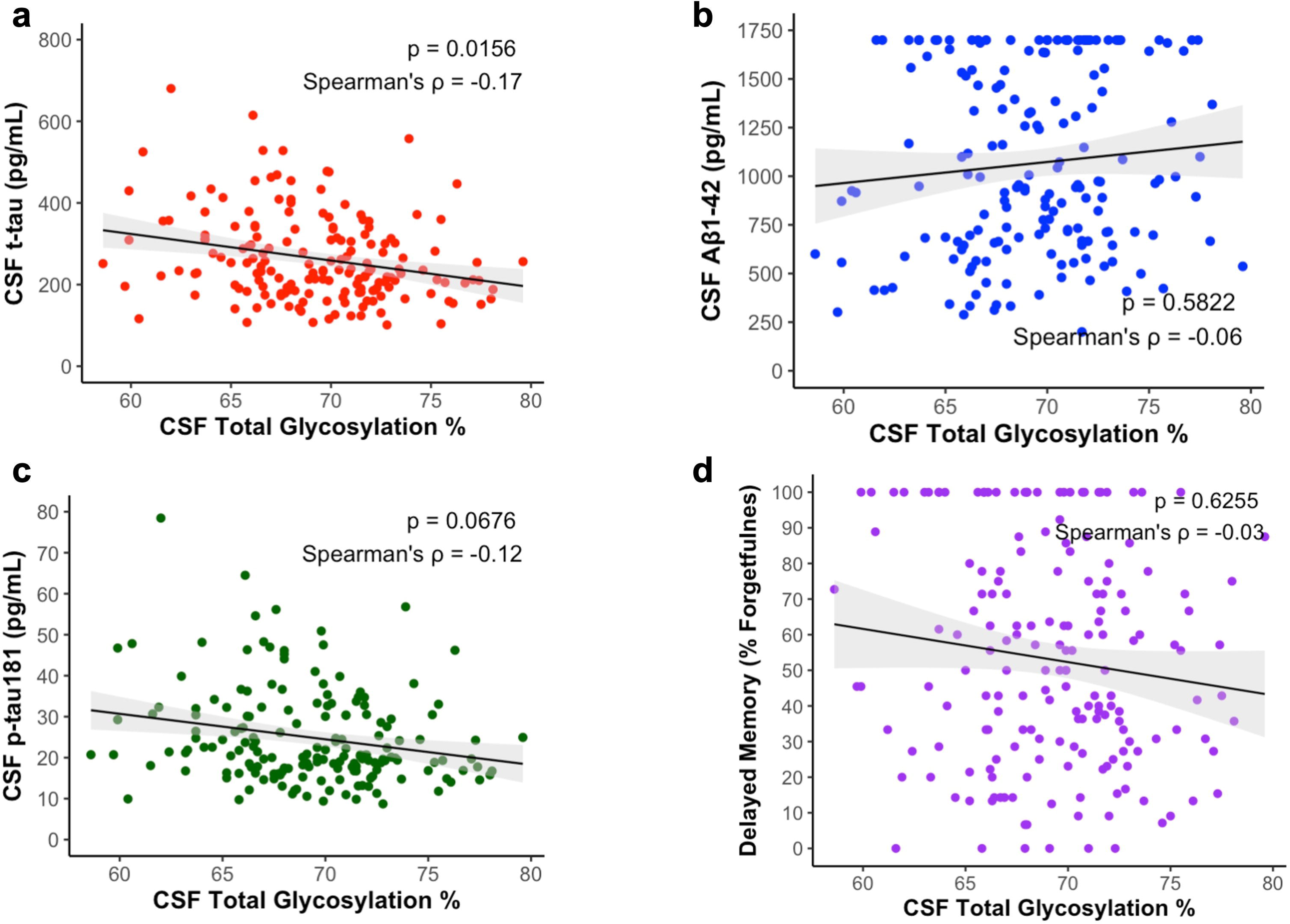
Baseline association of CSF total apoE glycosylation with CSF t-tau, p-tau181, Aβ_1-42_, and RAVLT % forgetting. Associations between apoE glycosylation and CSF AD biomarkers at baseline (t-tau, n = 174; p-tau181, n =; Aβ_1-42_, n =; RAVLT, n = 178) at baseline were examined using multiple linear regression models adjusted by age, sex, race, education, APOE4 genotype and clinical diagnosis.

#### APOE4 glycosylation is associated with lower tau pathology biomarkers, particularly in *APOE***ε**4/**ε**4 carriers

When analyzing the association of apoE3 and apoE4 glycosylation in the CSF with AD biomarkers, we found that apoE4 glycosylation (r = -0.34, p = 0.002), and less so apoE3 glycosylation (r = -0.16, p = 0.054), were associated with lower total-tau levels **(Fig 4a, b**). A similar pattern was observed for ptau-181, where apoE4 glycosylation correlated with lower ptau-181 levels in the CSF (r = -0.27, p = 0.012), with a weaker correlation with apoE3 glycosylation (r = -0.14, p = 0.101) (**Fig. 4c, d**). Interestingly, the negative association of apoE4 glycosylation with both total-tau (r = -0.57, p = 0.0001) and ptau-181 (r = -0.47, p = 0.009) were particularly stronger in *APOE*ε4/ε4 (**Fig. 4 b, d Supplement**). This effect was not observed for apoE3 glycosylation in *APOE*ε3/ε3 (**Fig. 4 a, c Supplement**).

**Fig 4.**
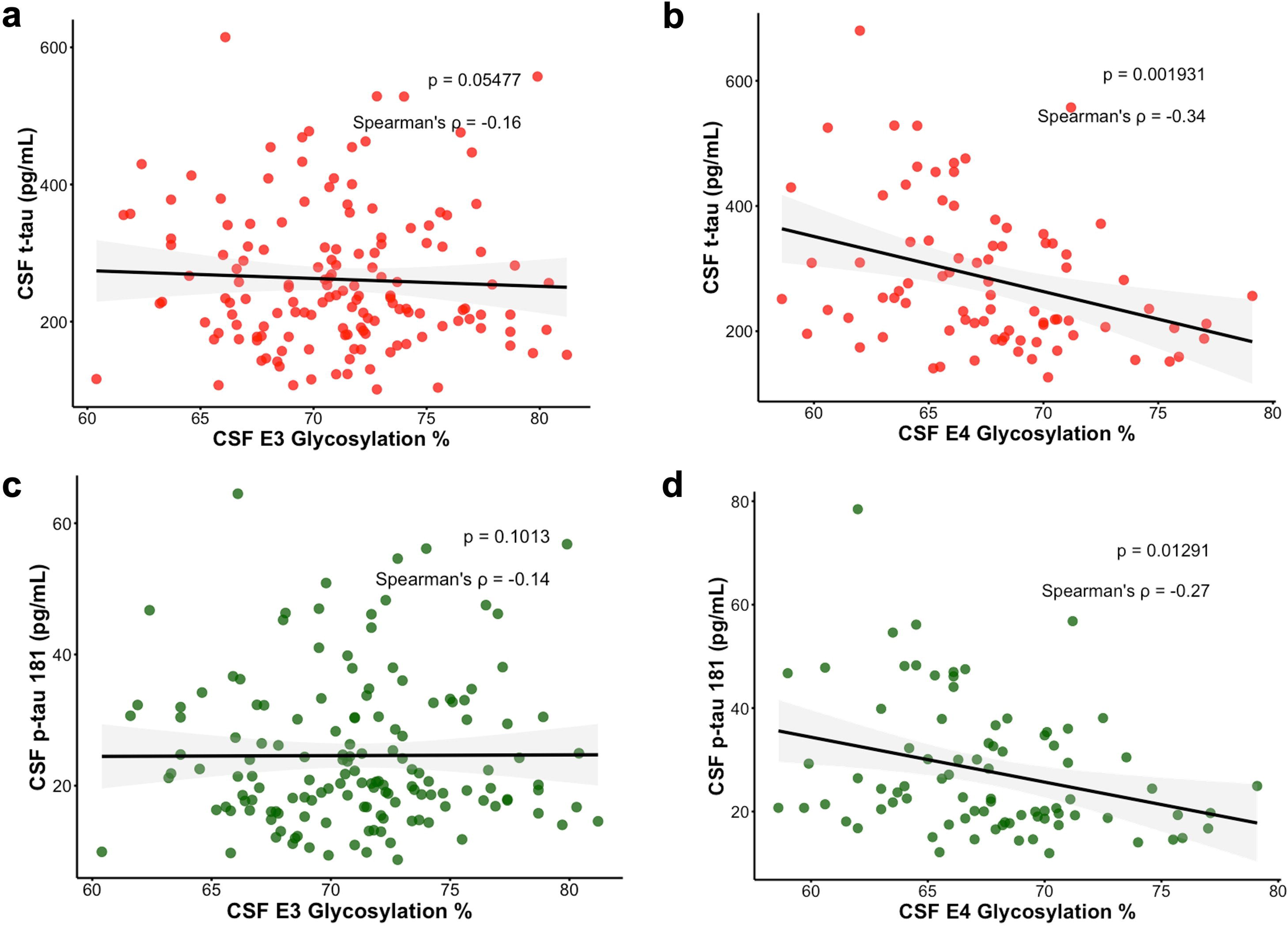
Baseline association of CSF total apoE3 and apoE4 glycosylation with CSF t-tau, p-tau181. Association of CSF apoE3 (A) and apoE4 (B) glycosylation with t-tau, and CSF apoE3 (C) and apoE4 (D) glycosylation with p-tau 181 levels in the CSF. The multiple linear regression models were adjusted by age, sex, race, education, *APOE*4 genotype and clinical diagnosis.

Regarding the effects of APOE secondary glycosylation in the CSF on tau pathology biomarkers, we observed no significant effects of either apoE3 or apoE4 secondary glycosylation on total-tau or ptau-181 levels (**Fig. 5 Supplement**). However, when stratifying the analysis by APOE isoforms, we found that apoE4 secondary glycosylation was associated with reduced total-tau (r = -0.41, p = 0.023) and ptau-181 (r = -0.31, p = 0.091) in APOEε4/ε4 (**Fig. 6b, d Supplement**). In contrast, no effect of apoE3 secondary glycosylation on total-tau or ptau-181 was observed in APOEε3/ε3 (**Fig. 6a, c Supplement**). Similar results were observed when we analyzed the effects of apoE3 and apoE4 glycosylation in plasma. No effect of both plasma apoE3 and apoE4 glycosylation were observed in total-tau and ptau-181 (**Fig. 7 Supplement**). However, when stratifying the analysis by APOE isoforms, we observed that plasma apoE4 glycosylation was associated with lower total-tau (r = -0.56, p=0.024) and lower ptau-181 (r = -0.52, p = 0.042) in *APOE*ε4/ε4 (**Fig. 8 b, d Supplement**). No effect of apoE3 glycosylation to total-tau and ptau-181 was observed in *APOE*ε3/ε3 (**Fig. 8 a, c Supplement**).

**Fig. 5.**
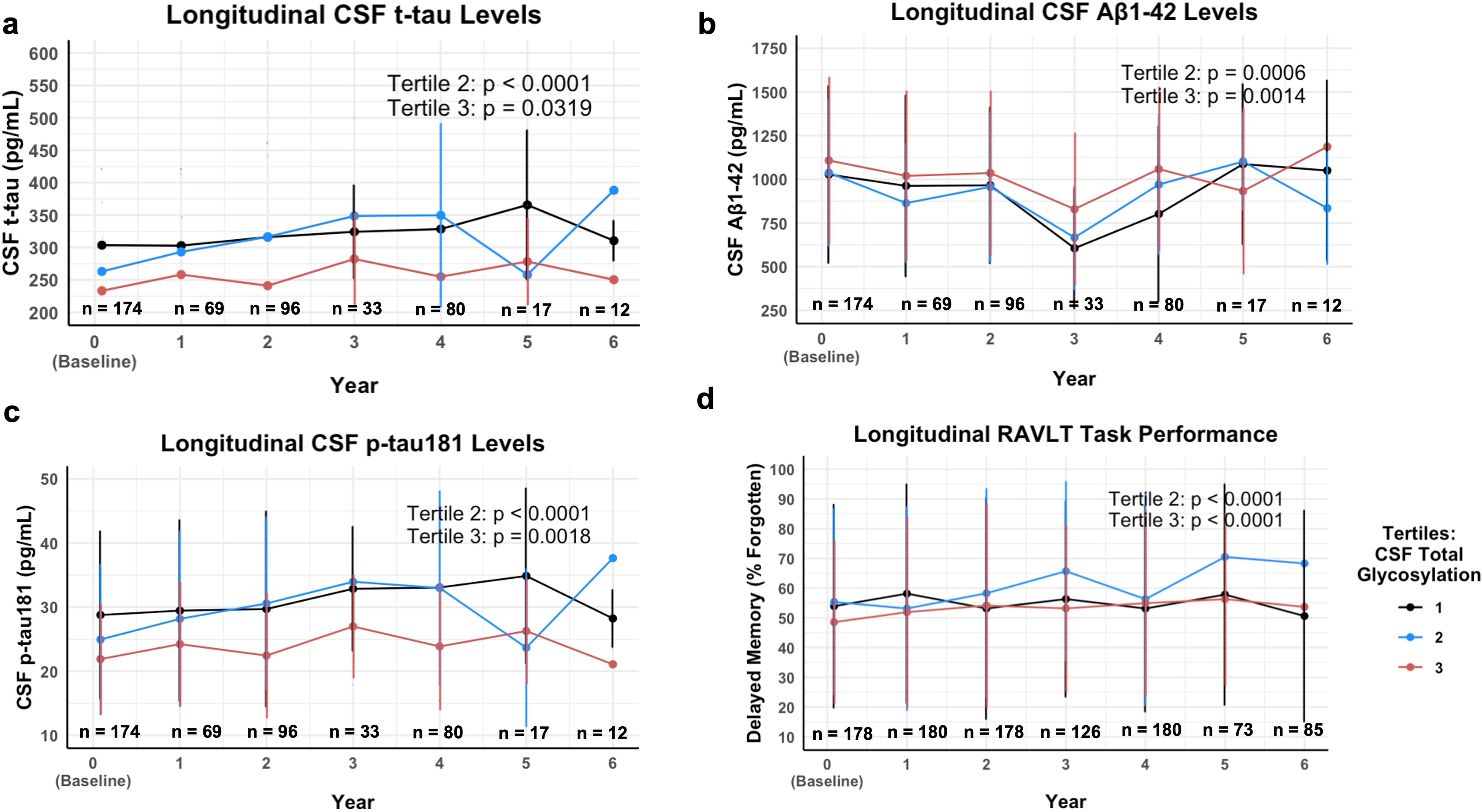
Longitudinal association of CSF total apoE glycosylation with CSF t-tau, p- tau181, Aβ_1-42_, and RAVLT % forgetting. ApoE glycosylation levels from the cohort (n = 181) were categorized into tertiles based on quantity (pg/mL). The associations between CSF apoE glycosylation tertiles and CSF AD biomarkers were examined using linear mixed effects models adjusted for age, sex, race, education, *APOE*4 genotype, clinical diagnosis, study site, and participant ID for repeated measures.

### Longitudinal analyses

#### Over a 6-year period, higher baseline levels of CSF apoE glycosylation predicted lower rates of increase in AD biomarkers

We used linear mixed effect models to evaluate whether apoE glycosylation tertile groups at baseline predicted longitudinal changes in AD biomarkers and RAVLT scores over six years of follow-up period. Cutoffs for tertiles of total glycosylation were ≤ 67.0 pg/mL (1^st^ tertile; “low glycosylation”); ≤ 71.0 pg/mL (“intermediate glycosylation”); ≤ 79.6 pg/mL (3^rd^ tertile, “high glycosylation”). Cutoffs for tertiles of secondary glycosylation were ≤ 16.5 pg/mL (1^st^ tertile; “low glycosylation”); ≤ 19.7 pg/mL (2^nd^ tertile “intermediate glycosylation”); ≤ 29.3 pg/mL (3^rd^ tertile; “high glycosylation”).

Intermediate levels of CSF total apoE glycosylation at baseline (compared to reference tertile 1) predicted greater CSF t-tau increase over time (p<0.0001), whereas having high glycosylation predicted significant CSF t-tau decrease over time (p=0.032, compared to reference tertile 1) (**Fig. 5a**). Intermediate levels of CSF total apoE glycosylation predicted greater increase in CSF p-tau181 longitudinally (p<0.0001 for the 2^nd^ tertile) while high glycosylation predicted significant CSF p-tau181 decrease over time (p=0.0018 compared to reference tertile 1) (**Fig. 5c**). Both intermediate and high levels of CSF total apoE glycosylation predicted lower CSF soluble Aβ_1-42_ decrease over time (p=0.00067 for intermediate glycosylation; p= 0.0015 for the high glycosylation) (**Fig. 5b**), and higher rate of increase in score on the RAVLT cognitive performance task (% forgotten) (p<0.0001 for intermediate glycosylation; p<0.0001 for high glycosylation) comparatively against reference tertile 1. (**Fig. 5d**). Similarly, high levels of CSF secondary apoE glycosylation at baseline (compared to reference tertile 1) predicted lower CSF t-tau increase over time (p=0.00012) (**Fig. 9a Supplement**), lower CSF ptau- 181 increase (p=0.00026 for intermediate glycosylation; p<0.0001 for high glycosylation) (**Fig. 9c Supplement**), and lower CSF soluble Aβ_1-42_ decrease over time (p<0.0001 for intermediate glycosylation; p=0.0009 for high glycosylation), (**Fig. 9b Supplement**). Intermediate levels of CSF secondary apoE glycosylation also predicted greater increase in % forgetfulness on the RAVLT cognitive performance task (p<0.0001 compared to reference tertile 1) (**Fig. 9d Supplement**). Greater levels of plasma total apoE glycosylation at baseline (comparatively against reference tertile 1) predicted higher CSF t-tau but not CSF p-tau181 increase over time (p<0.0001 for intermediate glycosylation, **Fig. 10 a, c Supplement**), lower CSF soluble Aβ_1-42_ decrease over time (p<0.0001 for intermediate glycosylation; p<0.0001 for high glycosylation, **Fig. 10 b Supplement**), and lower increase in % forgetfulness on the RAVLT cognitive performance task (p=0.011 for high glycosylation, **Fig. 10d Supplement**).

## Discussion

The apoE glycosylation results from the ADNI cohort presented in this work confirm that the three apoE isoforms have distinct glycosylation patterns, with *APOE*4 being the least glycosylated isoform in both CSF and plasma. These results are in close agreement with our two previous studies using different cohorts [27, 28]. ApoE secondary glycosylation was lower in the MCI group compared to the CN group and in progressors compared to non-progressors. At baseline, greater apoE glycosylation (total or secondary) was associated with lower CSF t-tau and p-tau181, regardless of *APOE* isoform. When grouped by protein isoform, CSF apoE4 but not apoE3 glycosylation was associated with lower t-tau and p-tau181 at baseline. In the longitudinal analysis, lower apoE glycosylation was associated with greater accumulation of CSF t-tau, CSF p-tau181, and lower CSF Aβ_1-42_ over a 6-year follow-up period, independent of *APOE* genotype. Overall, these results point to a protective role of apoE glycosylation on markers of AD progression, particularly on tau pathology.

This isoform-specific glycosylation trend has also been observed by several other groups. Moon *et al.* showed that apoE2 exhibits more sialic acid content than apoE3, with apoE4 being the least sialylated among the three isoforms [35]. Similarly, Mah *et al* showed that different apoE isoforms exhibit altered affinities for heparin, with apoE4 having the highest affinity, followed by apoE3, and then by apoE2 [36]. This heparin-binding apoE isoform trend is in close agreement with our previous results that showed increased binding of desialylated apoE to heparin [28]. Negatively charged sialic acids on the apoE glycan can repel the negatively charged heparin, reducing the binding of glycosylated/sialylated apoE. Thus, sialylation of apoE may play an important role in interactions with heparan sulfate proteoglycans (HSPG), which recognize apoE’s LDLR binding region and affect Aβ cellular uptake [37], promoting Aβ aggregation, glia cell inflammation response, and the propagation of toxic tau species.

The role of apoE glycosylation in AD pathology has not been established. However, glycosylation of other AD protein biomarkers has already been implicated. The amyloid-β precursor protein (APP) is glycosylated with an O-linked N- Acetylglucosamine-Galactose disaccharide (-GlcNAc-Gal) [38] and this glycosylation may inhibit Aβ production [39]. Tau, which is hyperphosphorylated in AD and forms neurofibrillary tangles [40], contains both N-linked and O-linked glycans: the N- glycosylation may be needed to induce hyperphosphorylation [41], while the O-linked glycosylation may act as a protective mechanism against hyperphosphorylation [42]. The enzyme glycoside hydrolase (O-GlcNAcase) removes the O-linked N- acetylglucosamine from tau, and there are several active drug development programs for inhibition of O-GlcNAcase to prevent the formation of neurofibrillary tangles [43]. However, the role and mechanism of apoE glycosylation in AD pathogenesis are likely more complex than those of tau. The apoE glycan is larger (GalNac(-Sia)-Gal-Sia) than the single GlcNac present on tau. Also, the presence of negatively charged sialic acids can change the biophysical properties of apoE more profoundly than the addition of a single GlcNAc on tau. It is very likely that the glycans on apoE (and their sialic acids) can modulate apoE interactions with its binding partners in vivo. Because CSF apoE is significantly more glycosylated than plasma apoE (and at two glycosylation sites in CSF vs. one in plasma), it is reasonable to assume that the apoE glycosylation is functionally more important in the brain. In one hypothesis, reduced glycosylation of apoE in brain could lead to reduced binding to Aβ and reduced ability to remove deposited Aβ plaques. In support of this hypothesis, mutant apoE3 with removed glycan attachment site exhibited reduced binding to Aβ_1-42_, and binding of fully glycosylated apoE with sialidase-removed sialic acids to Aβ_1-42_ was completely abolished [20]. Studies of Aβ have shown that binding to sialic acids reduces Aβ cell toxicity [44–46]. Furthermore, Lennol *et al.* recently observed that a less-sialylated form of apoE is present in higher proportion in AD subjects compared to controls [47]. In contrast, increased apoE binding to Aβ may decrease its clearance in mice [48]. Thus, less glycosylated apoE4 may increase Aβ accumulation due to reduced clearing. Then this in turn may increase tau phosphorylation. Indeed, increased tau load in humans is related to interaction between apoE4 and Aβ [49].

ApoE interactions or binding to tau [50], LRP1 [51], ABCA1 [52] and TREM2 [53] have already been demonstrated, but the role of the apoE glycan in these interactions is unknown. Of particular interest is the C-terminal region of apoE which contains the secondary glycosylation site at Ser290 (with possible sites also at Ther289 and Ser 296) [14, 26]. The C-terminal region of apoE presents a large exposed hydrophobic surface that initiates interactions with lipids, HSPG, and beta amyloid peptides [54], interactions which are most likely modulated by the attached glycan. The addition of glycans at this site may also increase the apoE solubility. Similar to the interaction between Aβ and apoE, changes in apoE binding to tau may directly or indirectly change tau dynamics. For example, glycosylation can further weaken apoE’s affinity for the microtubule-binding repeat region of tau, which is thought to be involved in the self-assembly of tau aggregates. ApoE3 is known to bind this region, likely helping to prevent tau aggregation and spreading [49]. Finally, N-glycosylation of tau is observed in the AD brain. Glycosylation of both tau and apoE may share some enzymes. If so, tau increasing in quantity in the AD brain may sequester these enzymes and prevent the proper glycosylation of apoE4. Or inversely apoE4 may hinder the function of these enzyme(s) reducing both tau and apoE glycosylation [55, 56].

We observed that apoE4 glycosylation, and less so apoE3 glycosylation, were associated with lower levels of tau pathology biomarkers. This finding is particularly significant given that the influence of *APOE* genotype on tau pathology is genotype-dependent [57]. The presence of the *APOE4* allele amplifies the effects of amyloid pathology on tau [49, 58, 59] and *APOE4* carriers have been shown to exhibit increased tau pathology biomarkers at early stages of the disease [57]. Building on these observations, we demonstrate for the first time that apoE4 glycosylation, but not apoE3 glycosylation, is negatively associated with tau pathology biomarkers, particularly in *APOE*ε4/ε4 carriers at early stages of AD. This effect also highlights the need to investigate the impact of apoE glycosylation on AD pathology in an APOE genotype-dependent manner. Although further mechanistic studies are needed to elucidate this effect, we propose that the post-translational addition of glycans to apoE4 increases its affinity for tau, thereby facilitating its clearance in the brain.

The study has some limitations that include the small size of samples in each of the four clinical groups, and the fact that the CSF and plasma samples were not matched (obtained from the same individuals). The selected ADNI samples did not represent diverse individuals or individuals with low educational backgrounds as education has an important role on cognitive performance. Results presented in this manuscript, need to be replicated and validated in larger, more diverse samples.

Furthermore, the three AD biomarkers were not measured in plasma samples, thus the plasma glycosylation data was only associated with CSF biomarker data for the same individuals at the same time points. Overall, these results are novel and point to a potential protective role of apoE4 glycosylation on brain tau pathology that merit further mechanistic studies.

## Acknowledgements

This work was funded in part by the National Institutes of Health/National Institute on Aging (NIH/NIA), RF1AG076124, R01AG055770, R01AG067063, R01AG054434, R21AG056518, and P30AG066530 to H.N.Y.; the Alzheimer’s Drug Discovery Foundation (ADDF) (GC-201711–2014197 to H.N.Y.), and donations from the Vranos and Tiny Foundations and Ms. Lynne Nauss to H.N.Y. M.S.R (AARFD-24-1313939) is supported by the Alzheimer’s Association Research Fellowship. T.A.P is supported by the NIH (#R01AG075336 and #R01AG073267) and the Alzheimer’s Association (#AACSF-20-648075). Data collection and sharing for this project was funded by the Alzheimer’s Disease Neuroimaging Initiative (ADNI) (National Institutes of Health Grant U01 AG024904) and DOD ADNI (Department of Defense award number W81XWH-12- 2-0012). ADNI is funded by the National Institute on Aging, the National Institute of Biomedical Imaging and Bioengineering, and through generous contributions from the following: AbbVie, Alzheimer’s Association; Alzheimer’s Drug Discovery Foundation; Araclon Biotech; BioClinica, Inc.; Biogen; Bristol-Myers Squibb Company; CereSpir, Inc.; Cogstate; Eisai Inc.; Elan Pharmaceuticals, Inc.; Eli Lilly and Company; EuroImmun; F. Hoffmann-La Roche Ltd and its affiliated company Genentech, Inc.; Fujirebio; GE Healthcare; IXICO Ltd.; Janssen Alzheimer Immunotherapy Research & Development, LLC.; Johnson & Johnson Pharmaceutical Research & Development LLC.; Lumosity; Lundbeck; Merck & Co., Inc.; Meso Scale Diagnostics, LLC.; NeuroRx Research; Neurotrack Technologies; Novartis Pharmaceuticals Corporation; Pfizer Inc.; Piramal Imaging; Servier; Takeda Pharmaceutical Company; and Transition Therapeutics. The Canadian Institutes of Health Research is providing funds to support ADNI clinical sites in Canada. Private sector contributions are facilitated by the Foundation for the National Institutes of Health (www.fnih.org). The grantee organization is the Northern California Institute for Research and Education, and the study is coordinated by the Alzheimer’s Therapeutic Research Institute at the University of Southern California. ADNI data are disseminated by the Laboratory for Neuro Imaging at the University of Southern California.

## Author Contributions

D.N. analyzed the samples and wrote the manuscript, Z.E.T. analyzed data and wrote the manuscript, M.S.R. analyzed data and reviewed the manuscript, S.D.H. helped design sample selection, I.S. provided analysis codes and study design, W.J.M. supervised the data analysis, T.A.P. reviewed the manuscript, H.N.Y. designed the study, obtained ADNI approvals, and wrote the manuscript.

