## Supplementary figures and images for "Increased cerebrospinal fluid and plasma apoE glycosylation is associated with reduced levels of Alzheimer’s disease biomarkers"

### Supplemental Fig 1

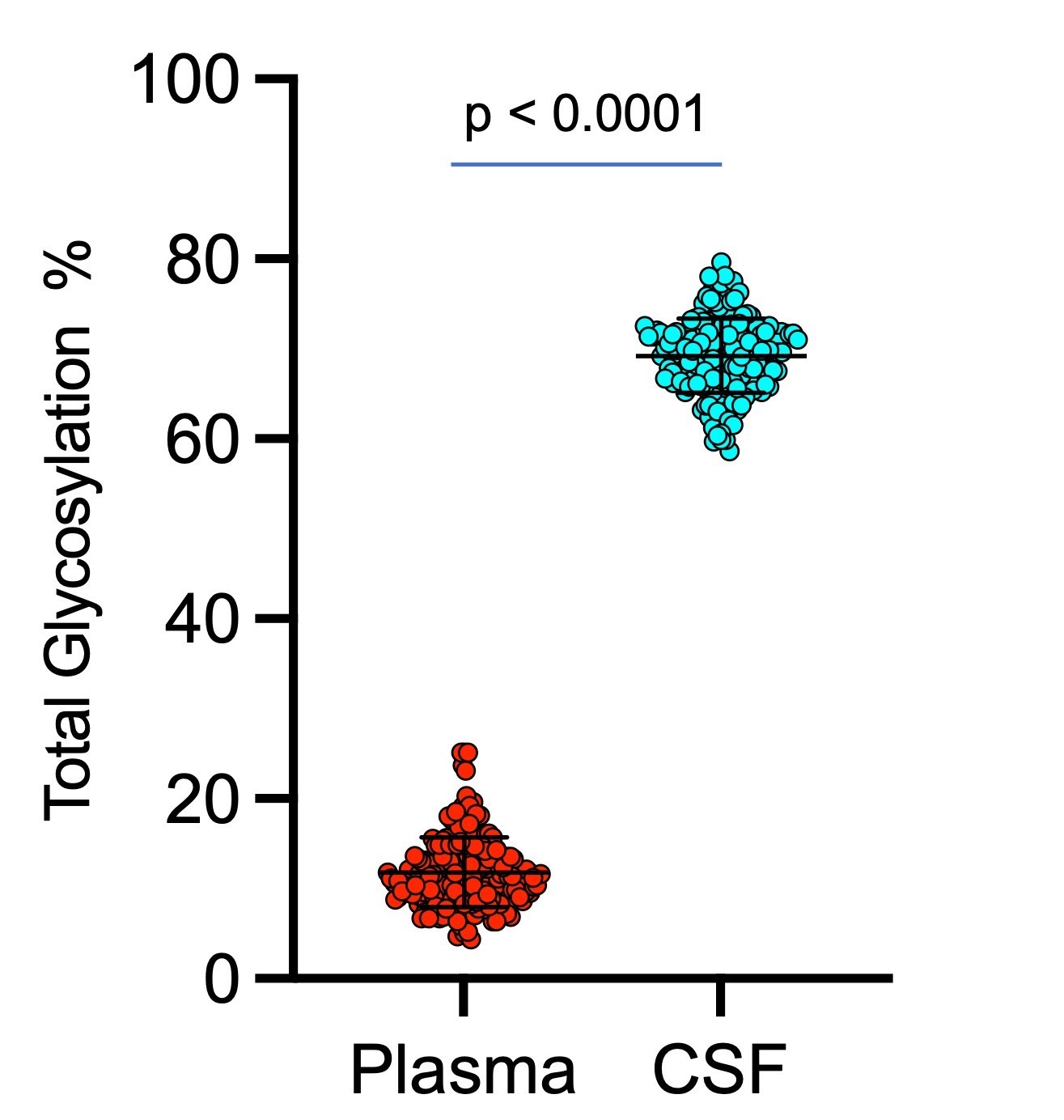

### Supplemental Fig 2

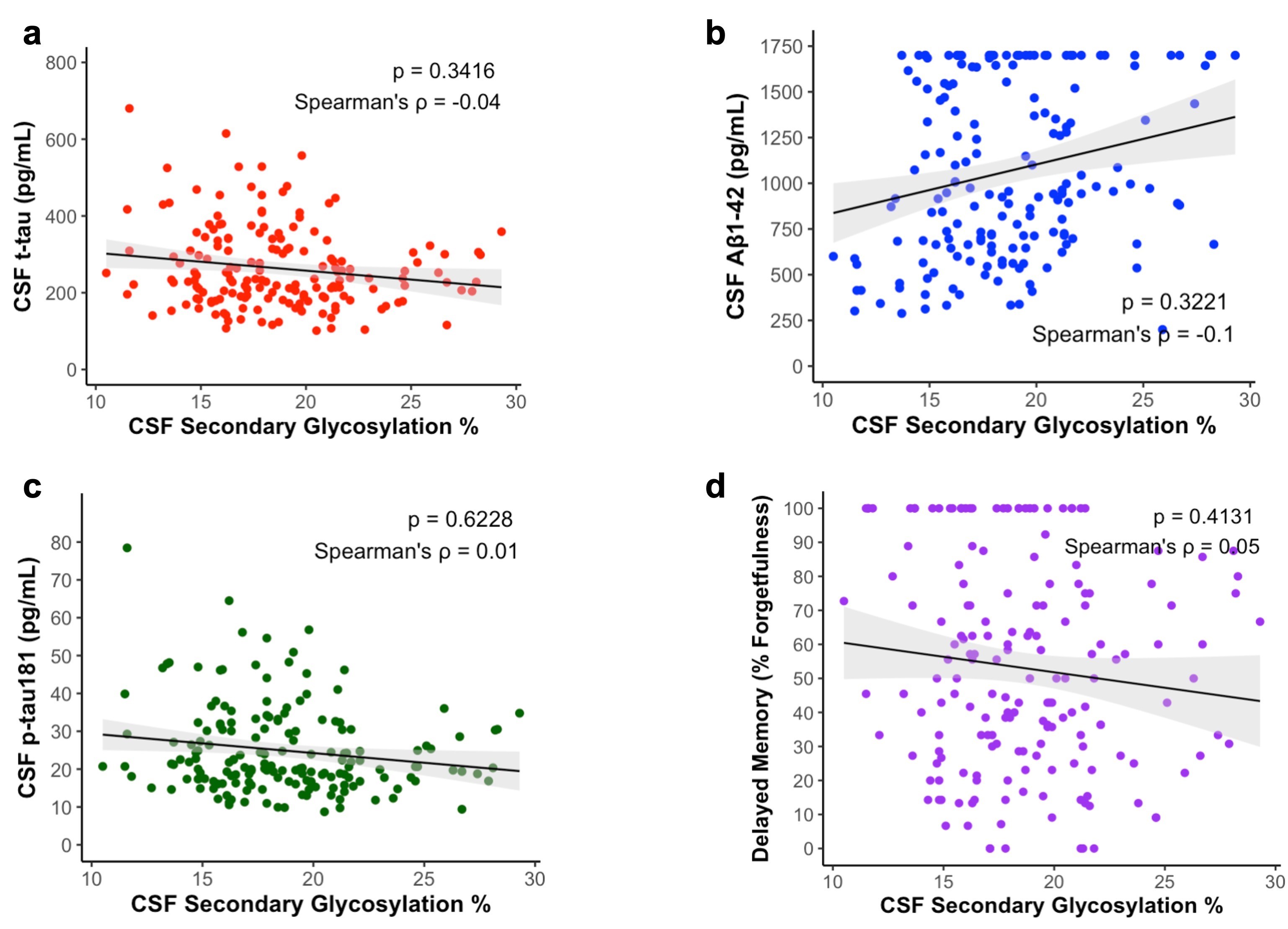

### Supplemental Fig 3

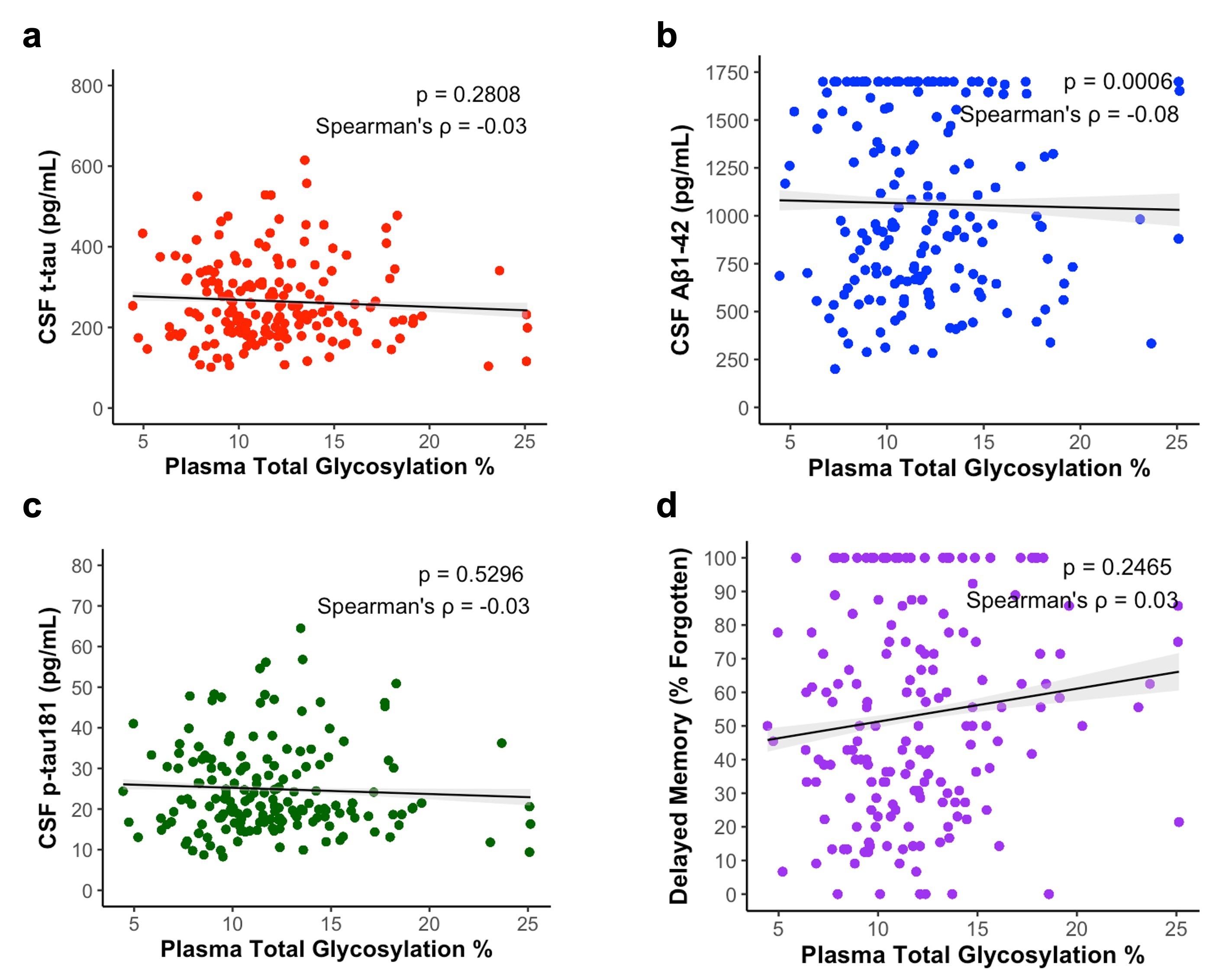

### Supplemental Fig 4

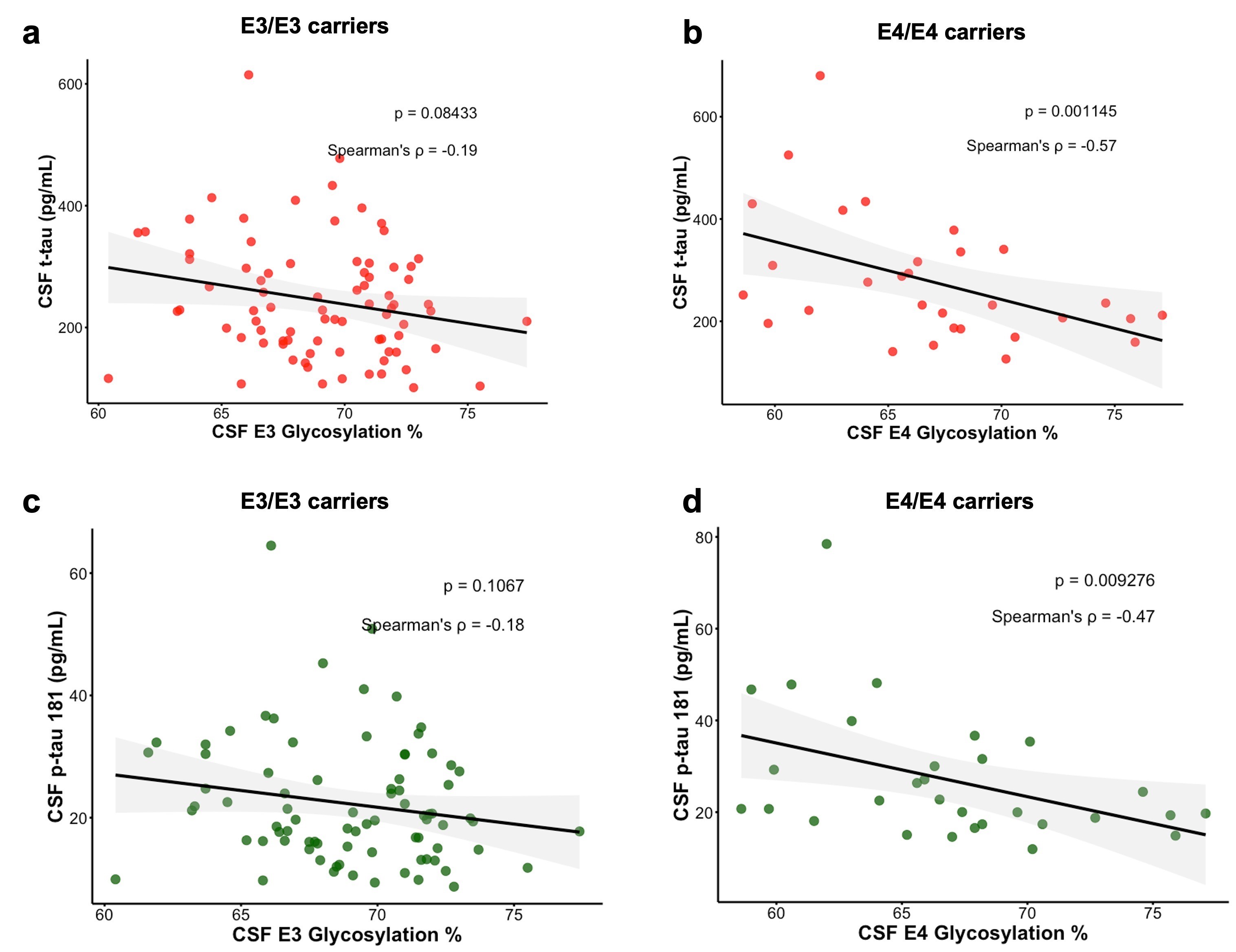

### Supplemental Fig 5

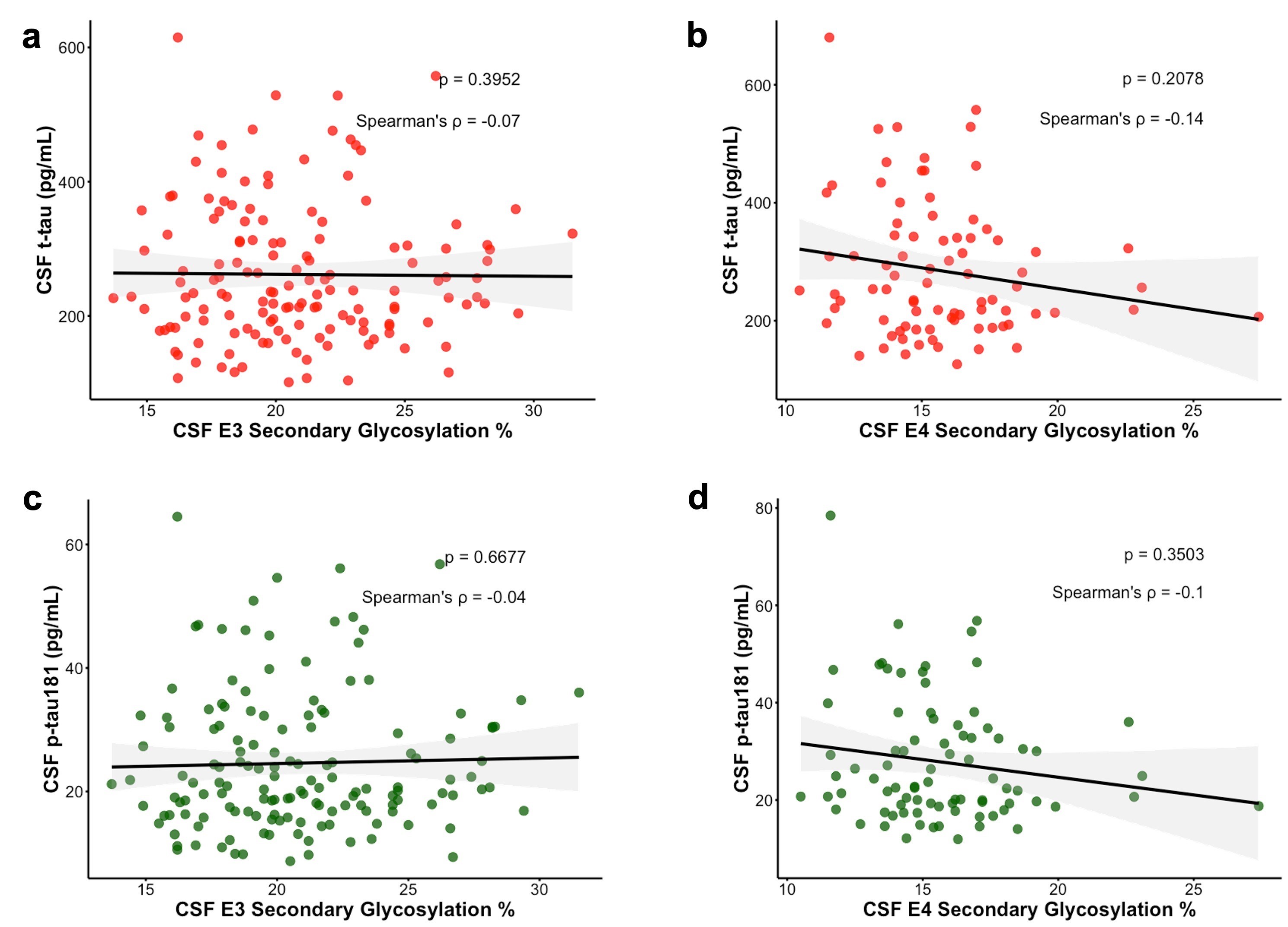

### Supplemental Fig 6

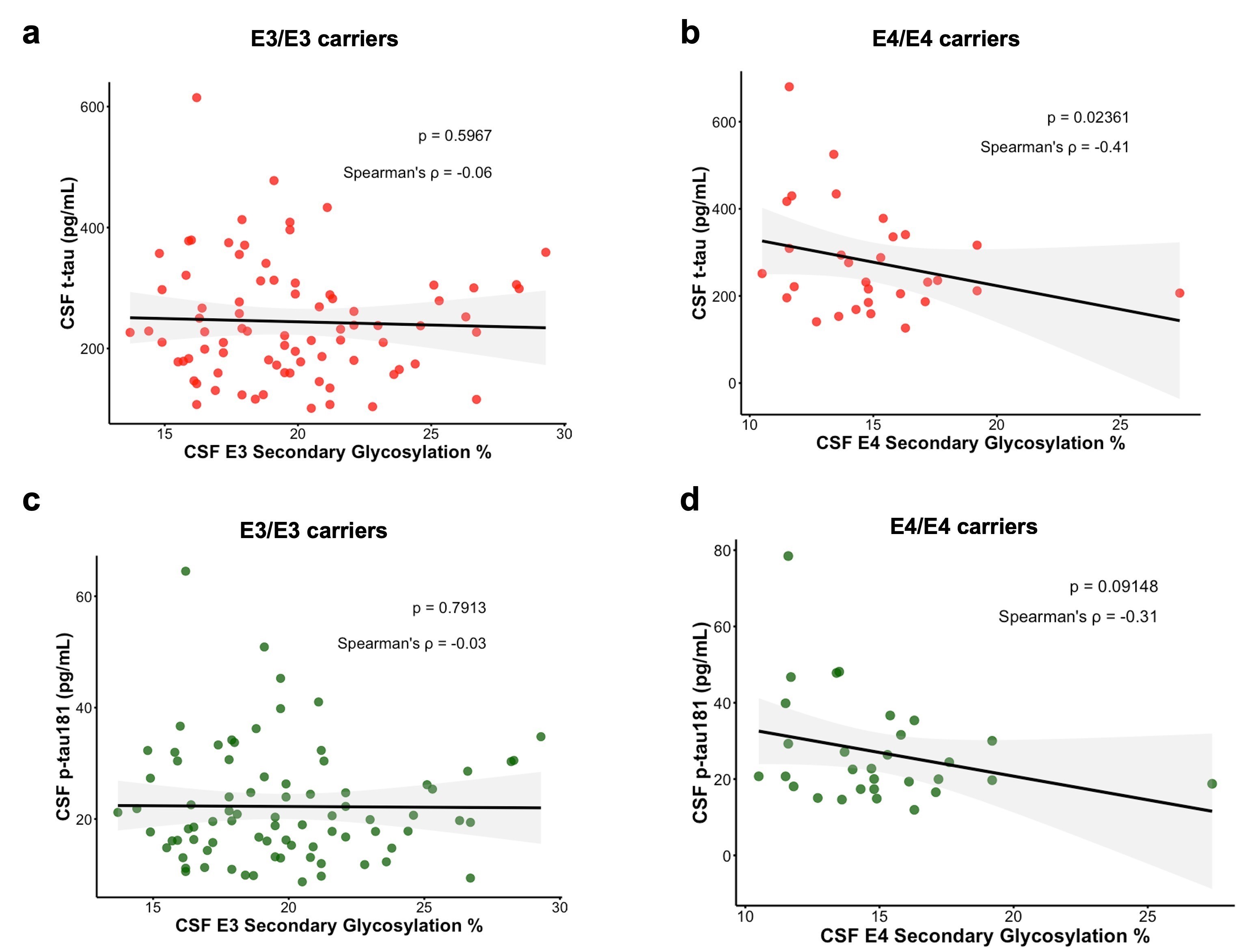

### Supplemental Fig 7

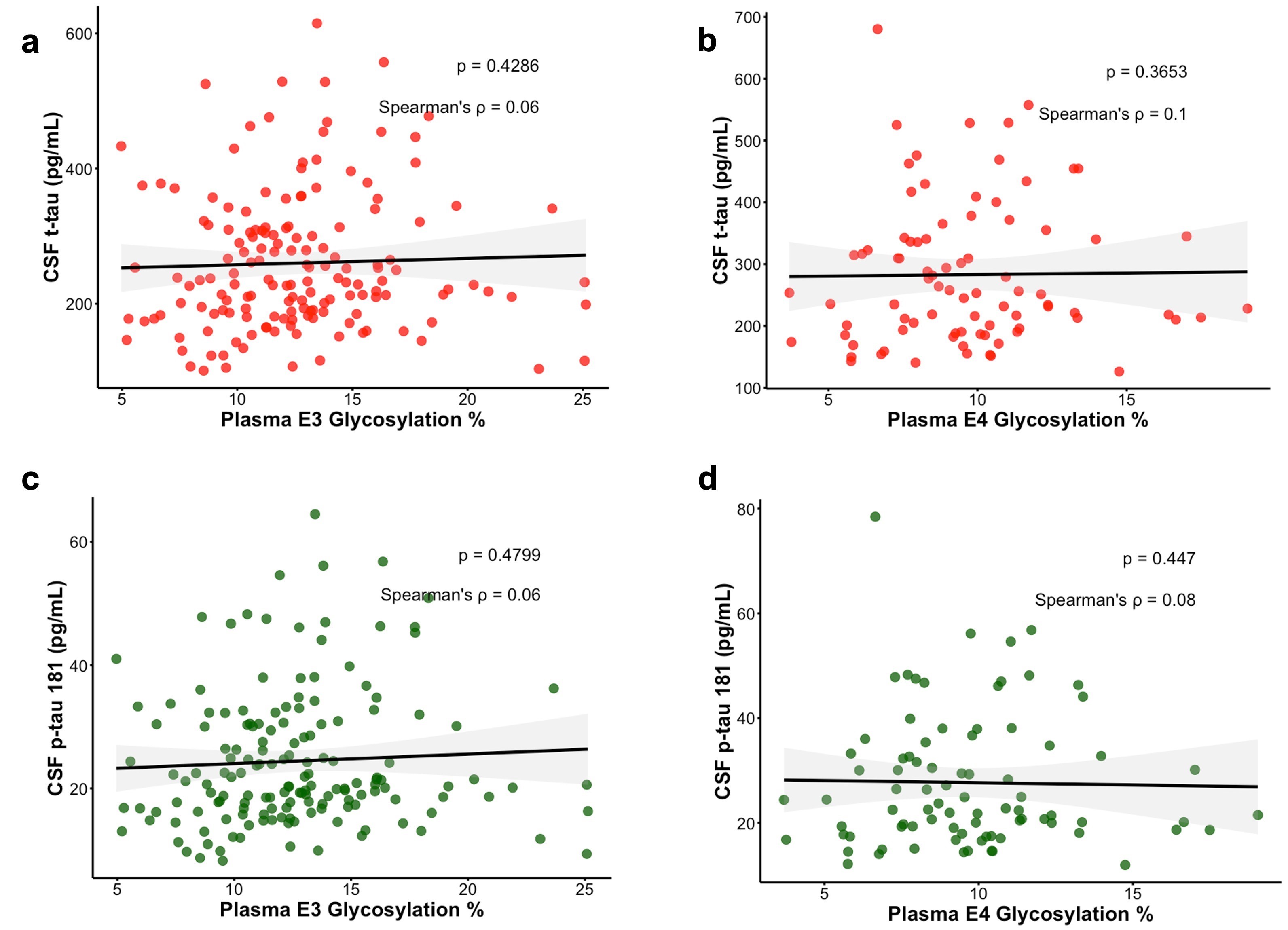

### Supplemental Fig 8

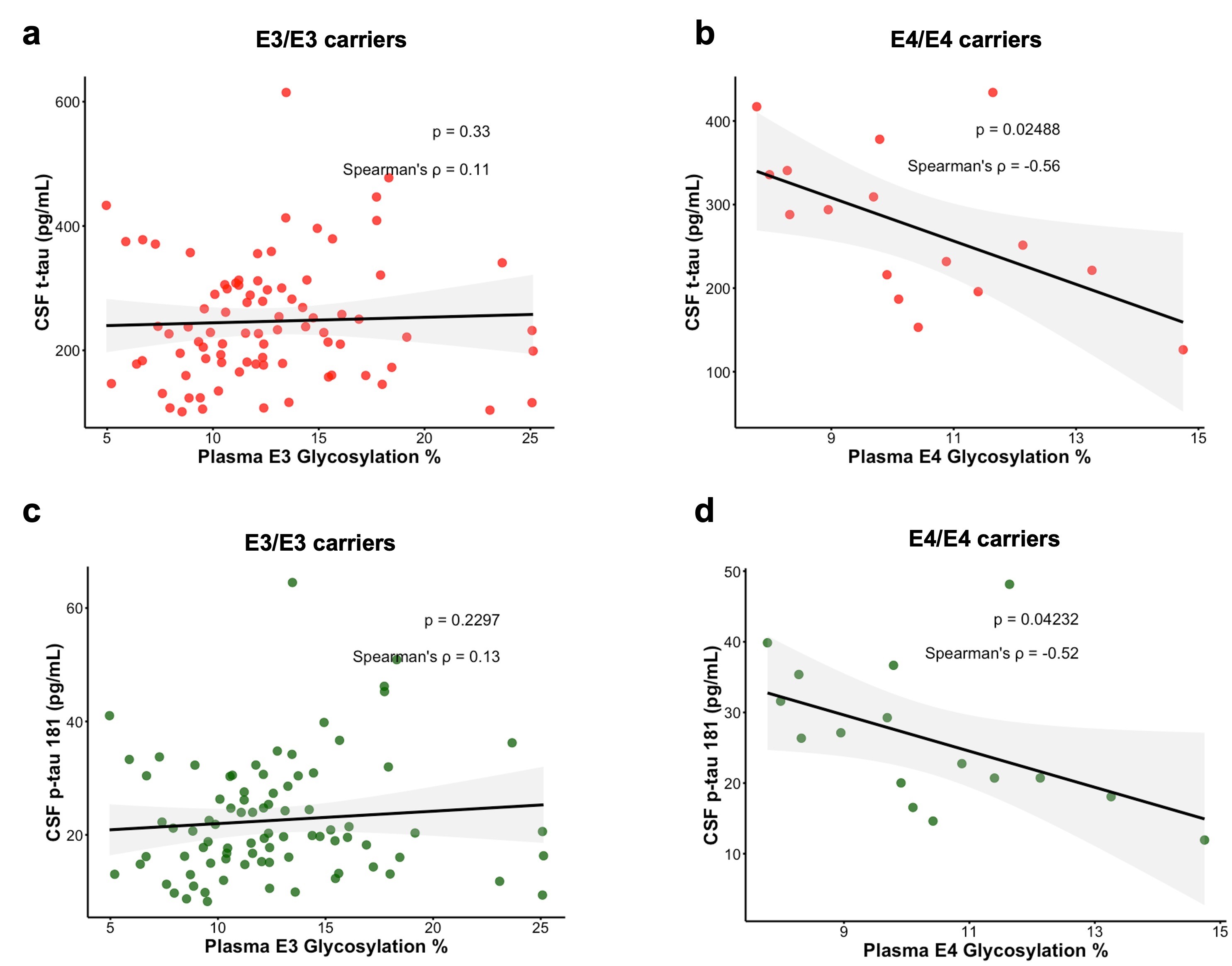

### Supplemental Fig 9

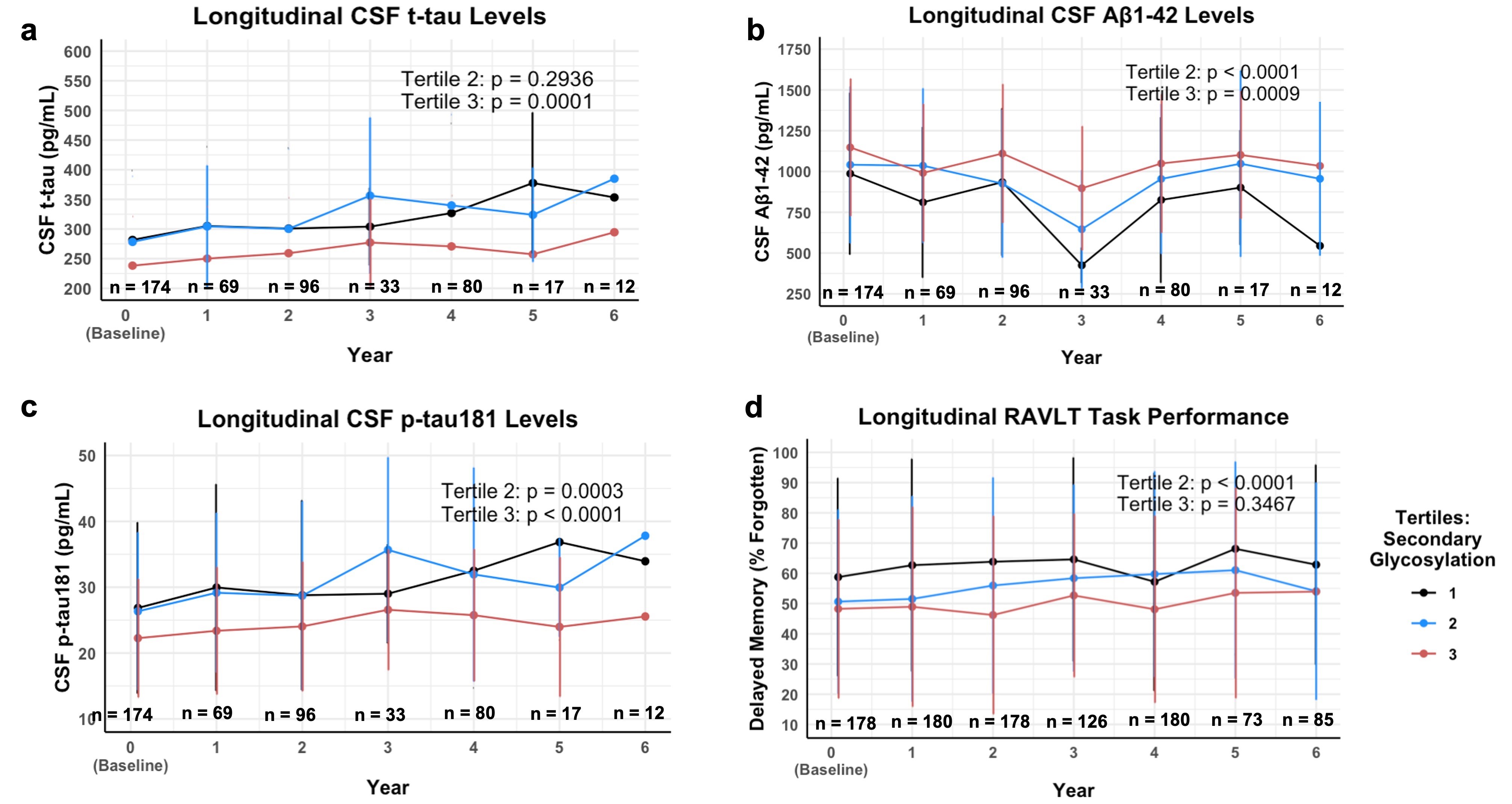

### Supplemental Fig 10

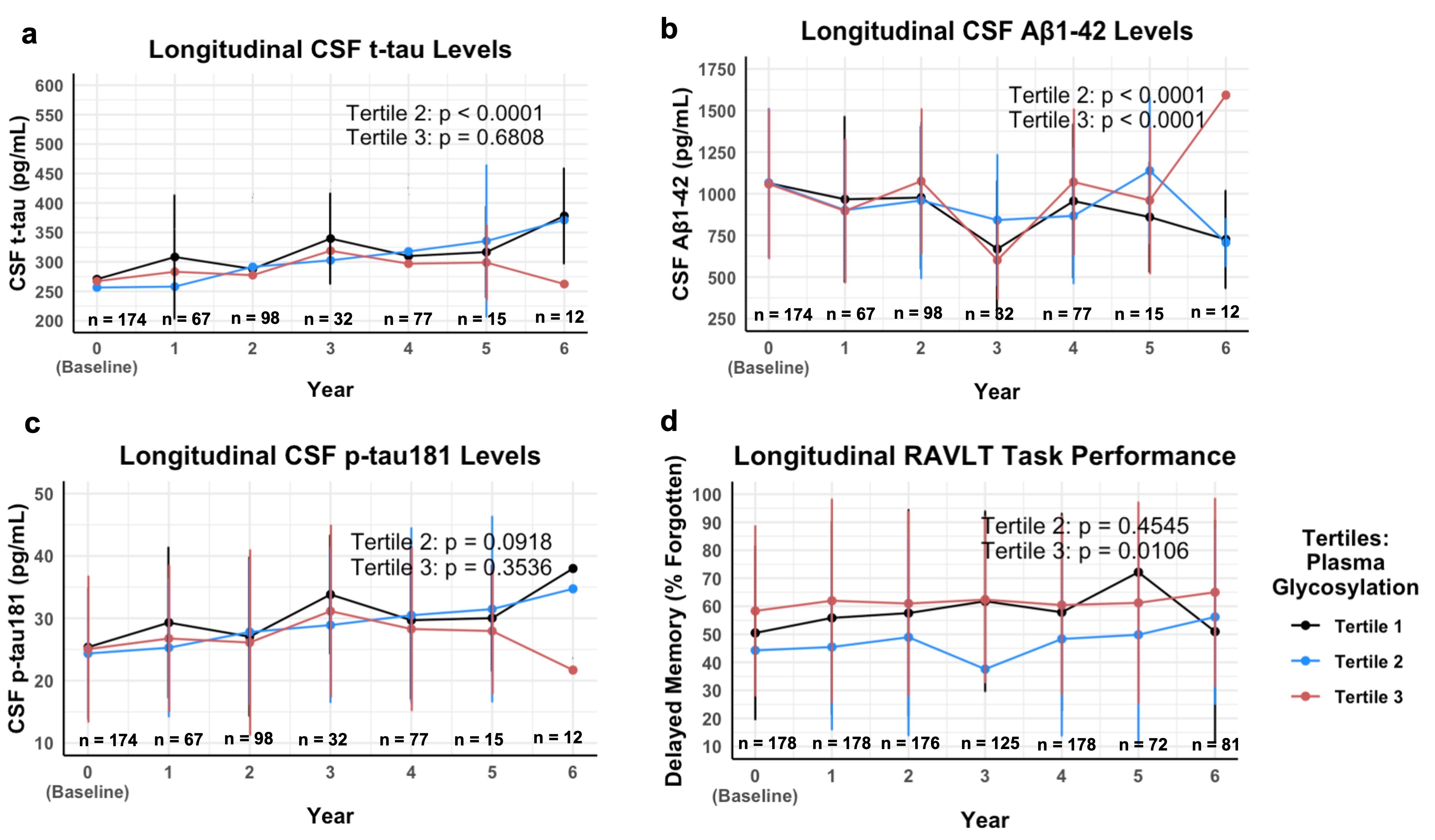
